# IL2 enhances ex-vivo expanded regulatory T cell persistence after adoptive transfer

**DOI:** 10.1101/805531

**Authors:** Scott N Furlan, Karnail Singh, Christina Lopez, Victor Tkachev, Daniel Hunt, James Hibbard, Kayla M Betz, Bruce R. Blazar, Cole Trapnell, Leslie S Kean

## Abstract

As regulatory T cell (Treg) adoptive therapy continues to develop clinically, there is a need to determine which immunomodulatory agents pair most compatibly with Tregs to enable persistence and stabilize suppressor function. Prior work has shown that mechanistic target of rapamycin (mTOR) inhibition can increase stability of thymic Tregs. In this study we investigated the transcriptomic signatures of ex-vivo expanded Tregs after adoptive transfer in the setting of clinically relevant immunosuppression using a non-human primate (NHP) model as a prelude to future transplant studies. Here, we found that adding interleukin-2 (IL2) to rapamycin *in vivo* supported a logarithmic increase in the half-life of adoptively transferred CFSE-labeled, autologous NHP Tregs, effectively doubling the number of cells in the peripheral blood Treg compartment compared to Treg infusion with rapamycin alone. Using single cell transcriptomics, we found that transferred ex-vivo expanded Tregs initially exhibit a gene expression signature consistent with an activated state. Moreover, those cells with the highest levels of activation also expressed genes associated with p53-mediated apoptosis. In contrast, transferred Tregs interrogated at Day +20 post-transfer demonstrated a gene signature more similar to published profiles of resting Tregs. Together, these preclinical data further support combining IL2 and rapamycin *in vivo* as adjunctive therapy for *ex-vivo* expanded adoptively transferred Tregs and suggest that the activation status of ex-vivo expanded Tregs is critical to their persistence.

## INTRODUCTION

There is a growing clinical need for an efficacious, suppressive cellular therapy for autoimmune diseases and transplantation. However, current globally immunosuppressive regimens are often associated with undesired off-target toxicities and can be antithetical to immune tolerance; calcineurin inhibitors being key examples of this paradox^1^. In contrast, suppressive cell-based therapies, including CD4+/CD25hi/FOXP3+ Tregs, promise fewer off-target effects and have been shown to induce immune tolerance in animal models^2, 3^. Substantial efforts are being made to establish the optimal strategy to sustain adoptively transferred polyclonal, CD4+/CD25hi/CD127lo thymically-derived Tregs in clinical trials^4–16^. Long-term and feasible clinical strategies will require that Tregs be paired with drug-based immunosuppressive agents already being used in the targeted patients, as even temporary cessation of these agents may put patients at risk for disease progression or recurrence.

A formidable challenge of *ex-vivo* expanded Treg therapy is ensuring their long-term persistence^4, 5, 14, 15, 17, 18^. The mTOR inhibitor, rapamycin has been associated with increasing frequency of endogenous murine thymic Tregs (tTregs)^19, 20^ and peripheral Tregs (pTregs)^21, 22^. Using a non-human primate (NHP) model of adoptively transferred *ex-vivo* expanded Tregs, we previously showed that systemic rapamycin (rapa) affords a modest prolongation in Treg persistence compared to the calcineurin inhibitor (CNI) – tacrolimus (t1/2 for rapa = 67.7 hours, versus 47.4 hours for tacrolimus)^15^, likely explained by Treg’s requirements for calcineurin-dependent IL2 production by non-Tregs as previously shown in rodent models^1^. Rapa also stabilizes the functional phenotype and gene expression profile of endogenous^16, 19, 20, 23^ and adoptively-transferred Tregs^15^. However, as monotherapy, rapa failed to promote long-term persistence of adoptively-transferred, ex-vivo expanded, autologous Treg^15^.

Interleukin-2 (IL2) is an attractive adjunctive therapy for a suppressive cellular therapy as it has a number of beneficial effects on both endogenous (non-transferred) tTregs and pTregs. Low-dose IL2 supports pTreg expansion in culture^24^ and the persistence of adoptively-transferred thymic Tregs used to reverse established chronic GVHD in mice^25^. At low doses in patients with chronic GVHD, IL2 expands the endogenous Treg compartment and has been shown to be therapeutically beneficial^26, 27^. When given as an immune complex with an anti-IL2 mAb, IL2 half-life is prolonged, similarly increasing the Treg compartment in mice^28, 29^. IL2 complexes also stabilize the expression of the Treg-lineage master transcription factor, FOXP3 in TGF-B-induced pTregs^30^. We hypothesized that ex-vivo expanded Tregs’ exposure to high IL2 concentrations may render them particularly sensitive to cytokine withdrawal-induced death^31^ (CWID) after adoptive transfer, a sensitivity that could be ameliorated with systemic IL2 therapy. Given proven IL2 and rapamycin (IL2+rapa) advantages in supporting endogenous Treg expansion in small animal models^32, 33^ and patients^34^, we tested IL2+rapa for its capacity to prolong the half-life of autologously-derived, ex-vivo-expanded Tregs after adoptive transfer in an outbred, NHP model and performed flow cytometry and single cell transcriptomics to explore underlying mechanisms and correlations with lifespan and Treg subset dynamics after transfer.

## MATERIALS/METHODS

For full details of the materials and methods used in this study, see Expanded Materials and Methods.

### Isolation and ex-vivo expansion of Tregs

CD4+/CD25hi/CD127lo putative Tregs, from autologous donors were flow-sorted from PBMC and expanded as previously described^15^. A full description of isolation and ex-vivo expansion methodology is found in the Expanded Materials and Methods.

### Infusion and tracking of CFSE labeled Tregs

Expanded Tregs recovered after cryopreservation were labeled with CFSE and infused as previously described^16^. A full description of infusion and tracking methodology is found in the Expanded Materials and Methods.

### Immunosuppression and IL2

Animals were treated with rapa daily by intramuscular injection which began two weeks before the Treg infusion, to ensure a steady state level had been achieved when Tregs were infused. The dose of rapa was adjusted to achieve a target trough level of 5-15 ng/ml. Aldesleukin (IL2) injections given at 1 million units/m^2^ were started 5 days before adoptive transfer until day 60 post-transfer. The rationale for the Day -5 start was to evaluate effect of IL2 on endogenous Treg compartment prior to infusion of expanded Tregs.

### Single cell RNA-seq library prep

Single cell libraries from sorted populations of Tregs and controls were generated using a UMI-based, droplet-partitioning platform (10X Genomics) and sequenced using a NextSeq 500 (Illumina). Details of the bioinformatic pipeline and datasets used in this work can be found in Expanded Materials and Methods.

## RESULTS

### Subcutaneous low-dose IL2 prolongs adoptively transferred Treg half-life and increases endogenous Treg numbers

Given the precedent for both rapamycin and IL2 supporting endogenous Tregs^19–22, 24–28, 32–34^ and our prior studies of rapamycin administration on adoptively-transferred Tregs in NHPs^15^, we tested whether adding daily subcutaneous low-dose IL2 (1×10^6^ IU/kg, as given to treat chronic GVHD patients) to rapamycin,^15^, would prolong the half-life of expanded Tregs (labeled with CFSE to distinguish transferred from endogenous cells). Generation, cryopreservation, thawing, labelling and administration of Tregs is schematized in **Figure 1a** and described in detail in Expanded Materials and Methods. The animals received Treg infusions ranging in cell number from 3 - 30×10^6^/kg recipient bodyweight (Supplementary Table 1), doses that have been tested to prevent acute GVHD in patients. Detectible peripheral blood CFSE+ Tregs were significantly higher in IL2+rapa experiments compared to historical rapa-only or no rapa experiments (n = 3 each group, using the same animals analyzed previously with rapa only)^15^. Compared to rapa-only, IL2+rapa increased transferred Treg half-life from 2.7 to 19.0 days, with an alpha decay of 2.0 (rapa) vs 10.3 (IL2+rapa) days, and a beta decay of 6.9 (rapa) vs 83.7 (IL2+rapa) days. (**Figure 1b**).

**Figure 1.**
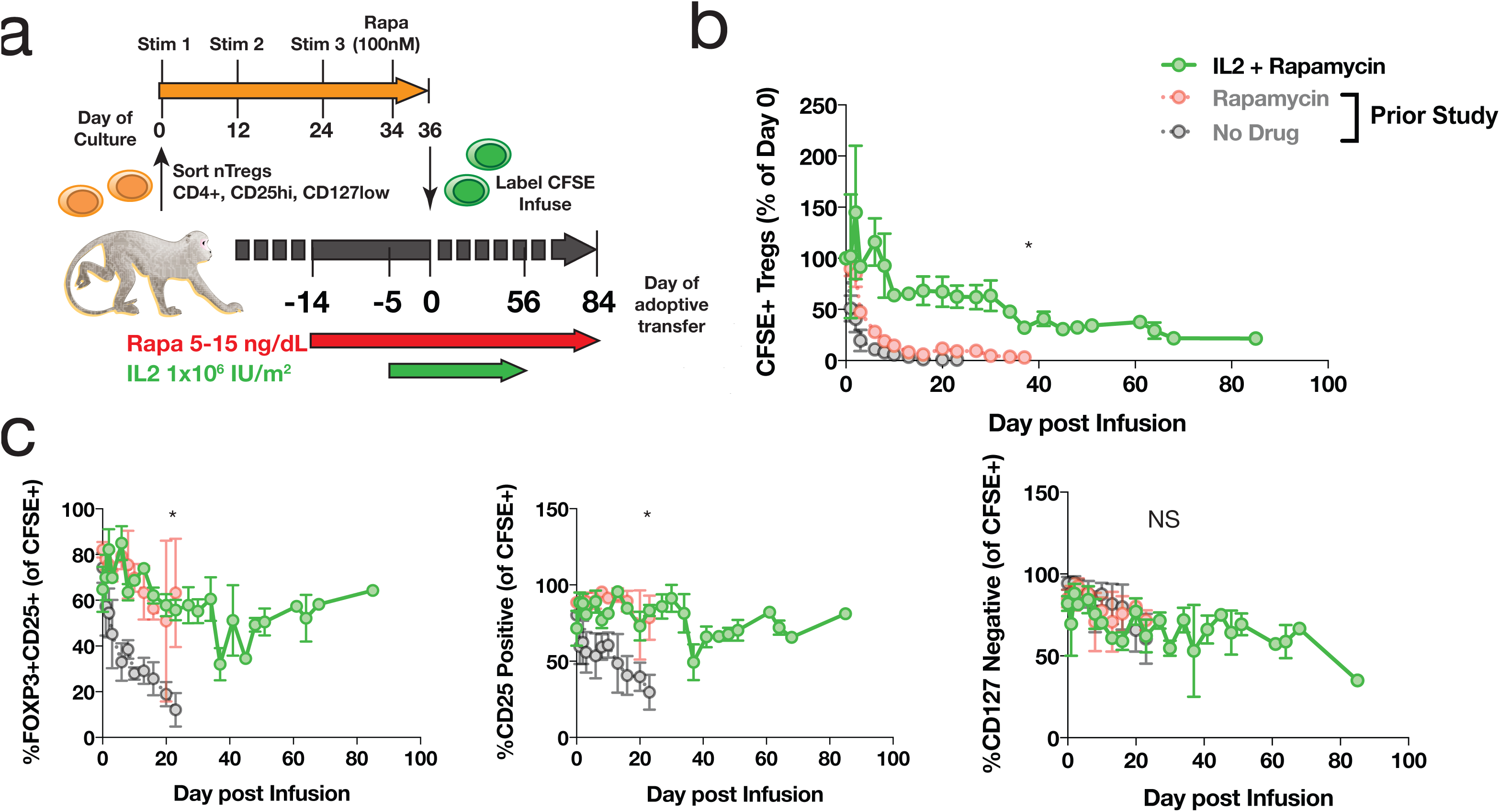
IL2 affords additional prolongation of T1/2 of adoptively transferred Tregs compared to rapamycin alone. a) Experimental schema. b) Longitudinal frequency of CFSE+ Tregs expressed as a percent of the 1 hour post infusion measurement. Tregs were defined as CD3+, CD4+, CD25+ and FOXP3+. * indicates P <0.05, for percentage of CFSE+ Tregs IL2+Rapa vs Rapa using repeated measures ANOVA. c) Percent of all CFSE+ cells that are positive for both CD25 and FOXP3 (left), CD25 only (middle) and negative for CD127 (right). * indicates P <0.05, comparing IL2+Rapa vs Rapa using repeated measures ANOVA, NS indicates not significant.

To ascertain the phenotypic characteristics of Tregs after IL2+rapa, levels of CD25, CD127 and FOXP3 expression in all CFSE+ cells ware assessed flow cytometrically in longitudinal fashion and compared to the rapa-only cohort. During IL2+rapa therapy, transferred Tregs retained expression of the canonical Treg phenotype (CD25hi/CD127lo/FOXP3+) to the same extent as previously observed with rapa alone (**Figure 1c**). As expected, IL2+rapa also resulted in a higher percent and absolute numbers of CFSE-negative endogenous (non-transferred) Tregs during IL2 administration in a time-dependent fashion (**Figure 2a-c**). Thus, when considering total cells in the Treg compartment, the combination of ex-vivo expanded Tregs with IL2+rapa, achieved a mean of 292 Tregs/uL blood, nearly tripling the mean counts of endogenous Tregs/uL blood (113 Tregs/uL) prior to Treg infusion. Longitudinal flow cytometric analysis of CFSE peaks, did not show multiple peaks of fluorescence as has been shown previously in NHP studies^16^ of actively dividing Tregs, nor did RNA-seq data (discussed later) reveal ongoing proliferation as evidenced by MKI67 or CCNB2 expression in the CFSE+ Tregs (**Supplementary Figure 2**).

**Figure 2.**
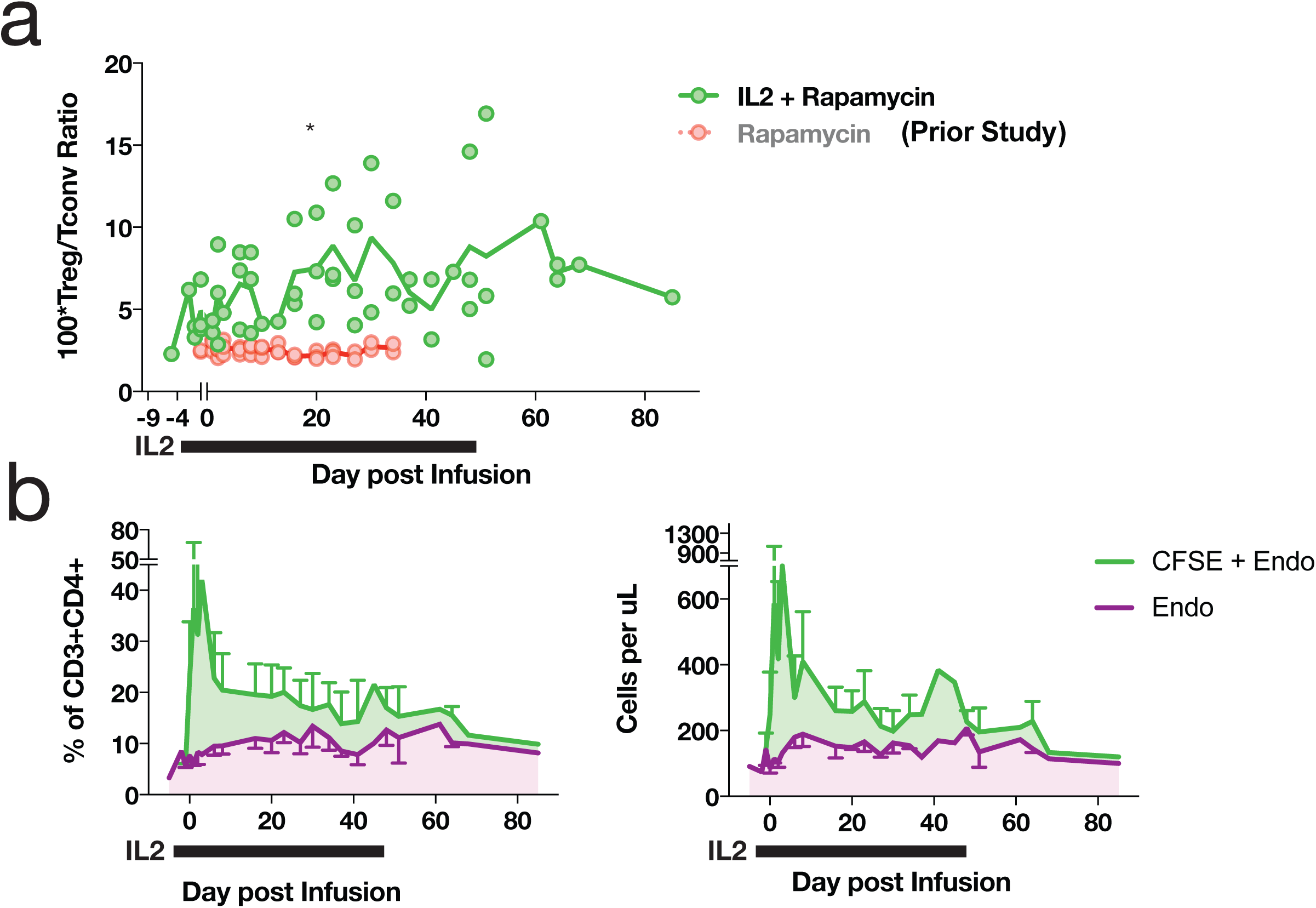
Subcutaneous low dose IL2 increases numbers of circulating endogenous Tregs. a) Treg/Tconv ratio x 100 in animals treated with IL2+rapa and rapa alone. * indicates P < 0.05 using repeated measures ANOVA b) Percentages (left panel) of circulating Tregs defined as a total percentage of CFSE+ cells (of CD3+CD4+) plus endogenous cells as gated in (a) - green; endogenous cells as gated in (a) alone - purple. Absolute numbers (right panel) of circulating tregs defined as an absolute number of CFSE+ cells (CD3+CD4+) per uL peripheral blood plus absolute endogenous cells as calculated in (a) - green; endogenous cells as calculated in (b) alone - purple.

### Single cell RNA-seq of endogenous and transferred Treg subsets

To both identify cellular heterogeneity in transferred Tregs and to identify gene expression patterns that might be associated with the persistence (or lack thereof) of transferred Tregs, we utilized single-cell Treg transcriptome profiling. To identify molecular events associated with the initial decline in transferred Tregs, cells were profiled at Day +3. To define those events associated with the prolonged persistence afforded by IL2+rapa we profiled again at Day +20. Importantly, studies to search for transcriptomic correlates with persistence were not technically feasible in prior experiments with rapa only due to the short half-life of transferred Tregs and as such, the improved half-life of Tregs with IL2+rapa allowed us to interrogate different phases of persistence of the adoptively transferred cells for the first time in our model system. To separately examine transferred Tregs, endogenous Treg and non-Treg control populations, we employed a flow cytometric sorting strategy that separated peripheral blood mononuclear cells into: 1) transferred Tregs (CD3+/CD4+/CFSE+); 2) ‘endogenous Tregs’ – (CFSE-/CD3+/CD4+/CD25h/iCD127lo); and 3) two control non-Treg populations of CFSE-/CD3+/CD4+/CD25hi/CD127hi or CFSE-/CD3+/CD4+/CD25lo/CD127hi cells (see **Supplementary Figure 1** for sorting strategy).

The sorting and scRNA-Seq analysis were performed on animal R.401, and sufficient and comparable numbers of cells from each population were obtained when sorting of PBMCs obtained at Day 3 and Day 20. Moreover, sorting ensured that single cell transcriptional profiles were generated from Tregs with validated Treg surface immunophenotypes. We recovered 25,037 single cell expression profiles from all timepoints that met quality control thresholds (see Expanded Materials and Methods) with a range of 1157-4751 cells per sorted cell type per timepoint.

We first performed analysis aimed at broadly visualizing clusters of gene expression profiles in specific sorted cell populations. Visualization utilized tSNE^35^ dimensionality reduction followed by DensityPeak^36^ clustering, an unsupervised clustering method used to define point density within a graph (**Figure 3a,b**). Distinct aggregates of sorted transferred and endogenous Tregs (upper left, blue and light-blue) and non-Tregs (upper right and lower middle [red and pink]) were observed (**Figure 3b**). Most clusters exhibited specific enrichment in sorted populations, supporting the notion that specific transcriptional differences underlie their canonical surface immunophenotype. In agreement with flow cytometric data, we identified canonical transcript expression levels associated with Tregs (high FOXP3, high IL2RA; low IL7R) in transferred cells (C1 and C2) and sorted endogenous Treg samples (C3 and C4, **Figure 3c**).

**Figure 3.**
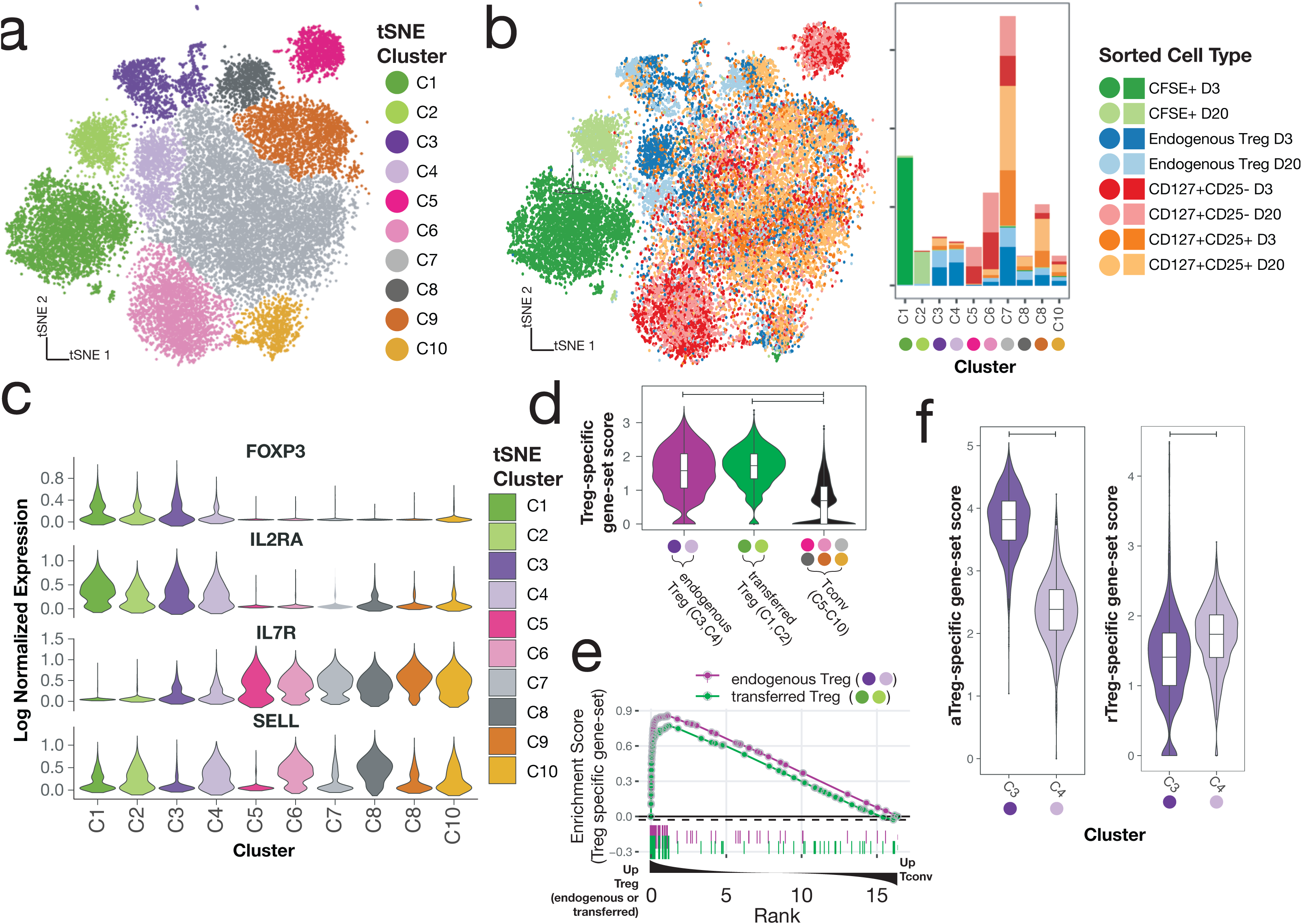
scRNA-seq reveals Treg subsets and shows that expression profiles of transferred Tregs maintain expression of Treg signatures after adoptive transfer. Visualization of single-cell expression profiles of all sorted cells after tSNE dimensionality reduction colored by unsupervised clustering analysis (a) and by sort gate (b-left panel) with yields of sorted cell types in each tSNE cluster (b-right panel). c) Expression of canonical Treg transcripts and SELL by tSNE cluster. d) Single cell scores of a Treg-specific gene-set signature taken from published profiles GSE90600 of human Tregs. e) GSEA enrichment plot of gene-set from (d) among differentially expressed genes between 1) endogenous Tregs and Tconv (tSNE clusters C5-C10) [green line] and 2) transferred Tregs and Tconv [purple line]. f) C3 and C4 tSNE clusters scored by human aTregs (left) and rTregs (right) gene-sets (GSE90600)

In addition to the clear clusters with the highest levels of FOXP3 expression (C1-C4), we observed that expression of CD62L+ (SELL) differentially marked clusters of both FOXP3-expressing and FOXP3-non-expressing cells (**Figure 3c**). Furthermore, we were able to note higher expression genes associated with T cell memory^37^ (CD58, CD63, LGALS3, S100A4) in most of the clusters cells that did not express SELL (**Supplementary Figure 3**), suggesting that naïve and memory T cell states, combined with the identity of a cell as either a Treg or not, are critical drivers of the gene expression programs in CD4 T cells. This paradigm is reminiscent of prior work describing five subgroups of cells in the CD4+ compartment^38^, namely: 1) “Fraction (Fr.) I”, aka “naive” or “resting” Tregs (herein ‘rTregs’) cells, distinguished as low levels of FOXP3 protein that are CD45RA^pos^; 2) “Fr. II” cells aka “activated” or “effector” Tregs (herein ‘aTregs’), distinguished by high levels of FOXP3 protein and that are CD45RA^neg^; 3) “Fr. III” cells commonly thought to be activated Tconv and exhibit low FOXP3 protein and are CD45RA^neg^ (herein ‘FrIII cells’); 4) CD45RA^neg^ non-Tregs (aka “memory” conventional T cells, herein ‘mTconv’); and 5) CD45RA^pos^ non-Tregs (aka “naïve” T cells, herein ‘nTconv’). Although CD45RA was not used in our sorting strategy, we hypothesized that the clustering patterns of our single cell expression data may be characterized by these previously described subtypes. Using previously published bulk transcriptomic profiles^39^ generated from flow-sorted populations of these 5 cell types, we set out to confirm the identity of our single cell cluster profiles.

As the cells in clusters C1 and C2 (transferred Tregs) and the cells in clusters C3 and C4 (endogenous Tregs) expressed the highest levels of FOXP3, we first sought to confirm that these cells exhibited a gene expression profile similar to both rTregs and aTregs as sorted Tregs in the aforementioned^39^ bulk transcriptomic dataset (GSE90600)^39^. To accomplish this using genes expressed in both resting and activated Tregs, we identified the intersect of the genes in the human GSE90600 dataset that were enriched in both: 1) rTregs compared to nTconv and 2) aTregs compared to mTconv. We then used this ‘Treg-specific’ gene-set to score each of our NHP single cell profiles (See Extended Materials and Methods for details of gene-set score calculation). Not surprisingly, clusters putatively identified as endogenous Tregs (C3 and C4) exhibited high expression for this gene-set (**Figure 3d**). Importantly, transferred Tregs (C1 and C2) also exhibited high expression of this gene-set to a level equivalent to that of endogenous Tregs (**Figure 3d**). To confirm these findings using another computational approach, we employed gene-set enrichment analysis (GSEA) to show enrichment of Treg-specific genes in both endogenous and transferred Tregs relative to Tconv (**Figure 3e**). As expected, canonical Treg markers, FOXP3, IL2RA, TNFRSF18, ENTPD1, CTLA4 were found in the leading edge of this genes enriched in both endogenous and transferred Tregs (**Supplementary Table 2** – GSEA leading edge genes). To classify cells in the C3 and C4 clusters, we used a similar approach of using gene-set scores (**Figure 3f**) which clearly identified the C3 cells as aTregs and the C4 cells as rTregs.

Although we identified that cells in C3 and C4 clusters were primarily cells from the flow-sorted ‘CD25hi/CD127lo’ population, many cells in this flow-sorted population did not cluster as bona-fide Tregs (**Figure 3b**) as is suggested by studies of CD25hi CD4 compartment heterogeneity^38, 39^. To further resolve all CD4+/CD25hi/CD127lo cells subsets using single cell RNA-seq, we then clustered only those cells sorted as endogenous Tregs (CFSE-/CD4+/CD25hi/CD127lo). From these 5690 cells, we identified 5 populations after dimensionality reduction and unsupervised clustering (**Supplementary Figure 3a**). Two clusters were easily identifiable as aTregs and rTregs being comprised chiefly of cells from cluster C3 and C4, respectively and having gene-set scores for published profiles^39^ consistent with their identity as such **(Supplementary Figure 3b)**. Another cluster comprised of cells chiefly from C8 and C9 original embedding, exhibited high gene-set scores for Fr.III cells (relative to aTregs) using published profiles^39^ while another cluster (Tconv) exhibited increased IL7R expression and decreased expression of IL2RA with absent FOXP3, and were derived from the original C7 cluster, supporting their identify as Tconv cells **(Supplementary Figure 3a-c)**. Finally, another cluster (herein referred to as ‘Other’) expressed FOXP3, IL2RA, and high levels of the STAT1 targets, ISG15 and MX1 (**Supplementary Figure 3d**). Consistent with this finding, these cells and the other C10 cells expressed high levels of a previously curated list of interferon gamma responsive genes^40^ (**Figure 3e**). While it is not immediately clear what this cluster of cells signifies, we interrogated an existing dataset of single cell RNAseq profiles in murine Tregs^41^ to evaluate whether these genes were observed in Tregs in other species. We obtained count data (GSE109742)^42^ and performed standard dimensionality reduction using tSNE (**Supplementary Figure 4**). We were able to clearly identify a small cluster of Tregs that express high Foxp3, Il2ra, Mx1, and Isg15 levels, arguing that this Treg subset is neither an artifact nor specific to the NHP. Further studies are ongoing toward a more complete characterization of this Tregs that exhibit downstream gene expression profiles consistent with STAT1-driven T cell activation.

Taken together, these results validate an approach that uses surface immunophenotype and single cell RNA-seq to identify circulating resting and activated Treg and non-Treg CD4 T cell populations. These data further build on prior work detailing human CD25hiCD4+ compartment heterogeneity^38, 39^ by confirming their existence in NHP Tregs and show that NHP resting and activated Tregs have distinct expression profiles resolvable by single cell RNAseq.

### Adoptively transferred Tregs gain transcriptomic similarity to endogenous resting Tregs with increasing time after transfer

Having rigorously characterized the cellular heterogeneity in the CD4+ compartment, we next set out to more precisely define the state of the transferred Tregs (clusters C1 and C2). As previously shown, these cells express a gene signature that is convincingly Treg in nature (**Figure 3d-e**). While there was a suggestion (based on expression of SELL) that the transferred cells at day 20 (cluster C2) may have a more “resting-like” profile than those at day 3, to evaluate this more definitively, we performed GSEA on C1 vs C2 cells using gene-sets derived from the endogenous NHP rTreg and aTregs as a reference (See **Supplementary Methods** for details on derivation of gene-sets). As shown in **Figure 4a (bottom)**, genes with increased expression in endogenous aTregs (C3) relative to their resting C4 counterparts exhibited clear enrichment in Day +3 transferred cells, while those with increased expression in endogenous rTregs (C4) exhibited a significant, but less complete enrichment at Day +20 (**Figure 4a - top**). As such, several transcripts enriched in C4 Tregs were more highly expressed in Day +3 cells (**Figure 4a – dashed ellipse**) suggesting that cells at Day +3 could be heterogeneous in resting- and activated-Treg gene expression. To interrogate this possibility further we used Uniform Manifold Approximation and Projection (UMAP), a manifold learning algorithm that optimally preserves global relationships in high dimensional data, to visualize the interrelatedness of transferred cells with cells from only the bona fide resting- and activated-Treg clusters (C3 and C4). This approach revealed that Day +3 transferred cells exhibited a far greater degree of transcriptional heterogeneity than Day +20 transferred cells. Unsupervised louvain clustering revealed 5 self-aggregating clusters of transferred cells. (**Figure 4b**). As suggested by the GSEA analysis, Day 20 transferred cells had high rTreg gene expression levels and clustered intimately with endogenous rTregs (**Figure 4b,c**). Day 3 cells separated into an “activated-like cluster” that expressed the highest aTreg gene levels relative to all other transferred cells, and an “intermediate” cluster that contained cells spanning UMAP space between the activated-like cluster and the more resting-like day 20 transferred cells. A small number of cells from both day 3 and day 20 comprised 2 other clusters (**Figure 4d**): 1) a cluster expressing high levels of the STAT1 target genes ISG15 and MX1 (similar to the STAT1-target expressing endogenous Treg cluster discussed previously) and, 2) a cluster of cells expressing high transcripts levels for cytolytic proteins (GNLY, CGA1, GZMB). Together, these data reveal vast heterogeneity in expression of resting- and activated-Treg genes in transferred cells early after adoptive transfer and highlight a more homogeneous expression of resting Treg genes seen by day 20.

**Figure 4.**
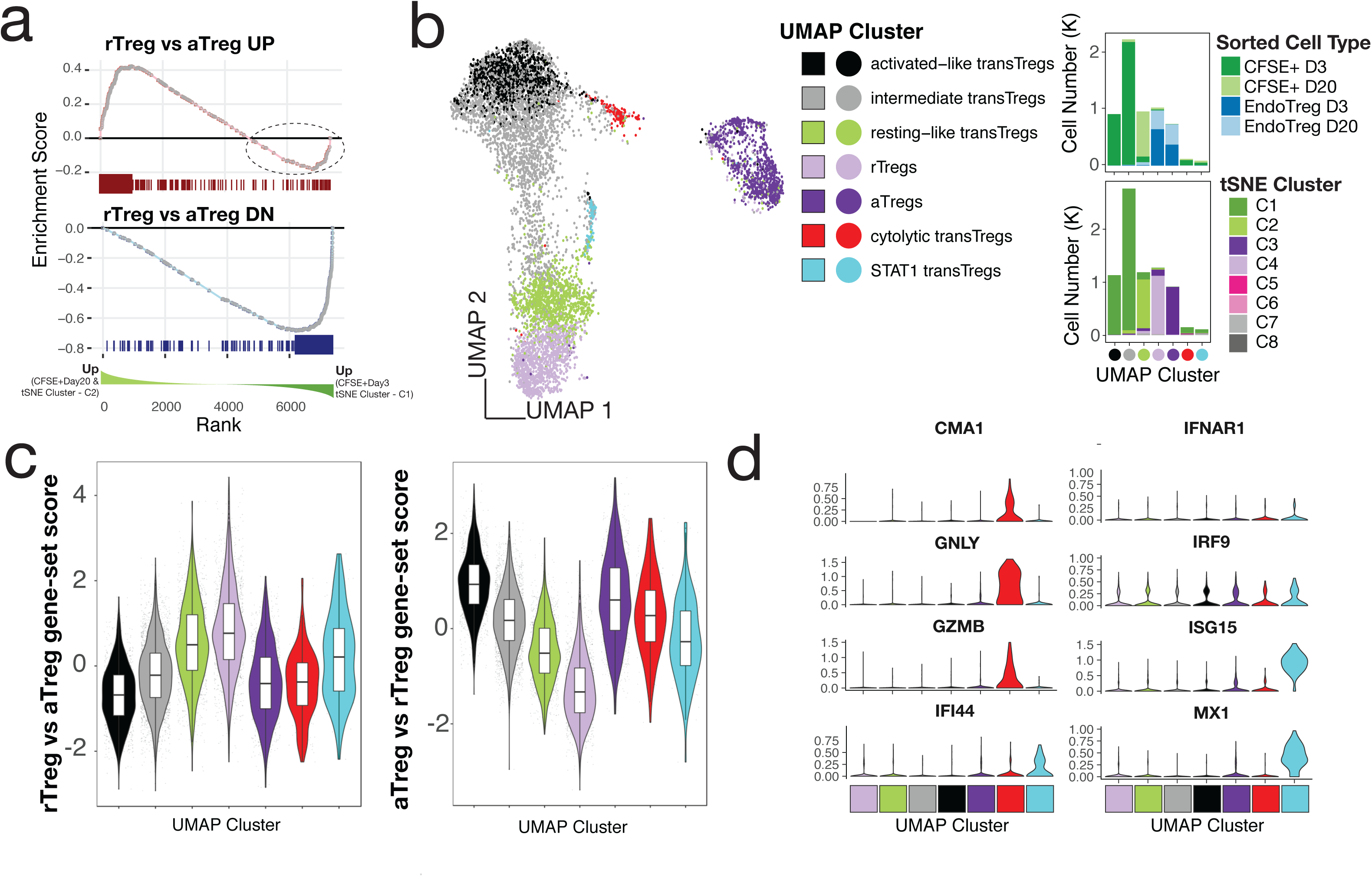
Transferred Tregs exhibit cellular heterogeneity and express a continuum of resting- and activated-Treg-related genes early after transfer, but are more “resting-like” by day 20. a-top) GSEA enrichment plot of NHP rTreg-specific genes in the ranked list of DE genes between Tregs at day 3 (tSNE cluster C1) compared to transTregs at day 20 (tSNE cluster C2). Dotted circle corresponds to those genes up in rTregs whose expression was higher in day 2 transferred cells compared to day 20. a-bottom) GSEA enrichment plot as in (a), but using NHP aTreg-specific genes. b-left) UMAP dimensionality reduction and clustering of all CFSE+ cells and cells from C3 and C4 tSNE clusters that were sorted as endoTregs. b - right) Cell percentages of sorted cell type (top) and tSNE cluster (bottom) in UMAP clusters from (a). c) Gene-set scores from (a) plotted in UMAP clusters (left) and UMAP embedding (right). d) Genes exhibiting high specificity scores for red and cyan clusters.

### Activated-like transferred Tregs exhibit profound p53-target gene enrichment and express high apoptotic gene levels

While the gene expression profile of Day +3 transferred cells was more activated than the Day +20 cells, the nonunion of the highly activated Day +3 cells with endogenous aTregs suggested there were significant underlying differences between Day +3 activated cells and endogenous aTregs (**Figure 4b**). To identify the gene expression changes that set these two groups apart, we performed differential expression on the activated-like transferred Tregs and endogenous aTregs (black vs purple cells– **Figure 4d**). Pathway analysis revealed that a significant number of genes overexpressed in activated-like transferred Tregs compared to endogenous aTregs were p53 pathway members (**Figure 5a-b**). Only 4 genes from this analysis were found to have decreased expression in activated-like transferred Tregs (TCHH, RPL36, S100A4 and RPS12). Of these, 3 (RPL36, S100A4 and RPS12) have been associated with a proliferative effect. Inhibition of RPS12 has previously been shown to function in concert with S100A4 to limit cell proliferation in cell lines^43^ and RPL36 has been associated with glioma cell proliferation^44^. Decreased expression of these proliferative genes in activated-like transferred Tregs suggest one possible mechanism for their disappearance over time. To confirm that p53 target genes, in addition to p53 pathway members, were increased in the activated-like transferred T cells, we utilized a consensus gene list from a recent meta-analysis^45^ of p53 target studies to calculate a ‘p53 target’ gene-set score for each cell. Scores were highest in Day +3 activated-like transferred Tregs relative to all other groups of transferred or endogenous cells. Because p53 has a diverse array of biologic functions, we sought to characterize the predominant p53 function in these cells by examining the top differentially expressed p53 targets (vs endogenous aTregs) and annotated them as Apoptotic, Anti-Apoptotic, Anti-proliferative, Proliferative, or Other. At a 2-fold change cutoff, no genes over-represented in activated-like transferred Tregs were either Proliferative or Anti-apoptotic, while 6 of the 11 genes favored apoptosis or were anti-proliferative (**Figure 5c**). These results suggest that early after adoptive transfer, transferred Tregs express a spectrum of activation states, the more activated of which is associated with an apoptotic gene expression profile, while the less activated (and less apoptotic) is likely associated with persistence.

**Figure 5.**
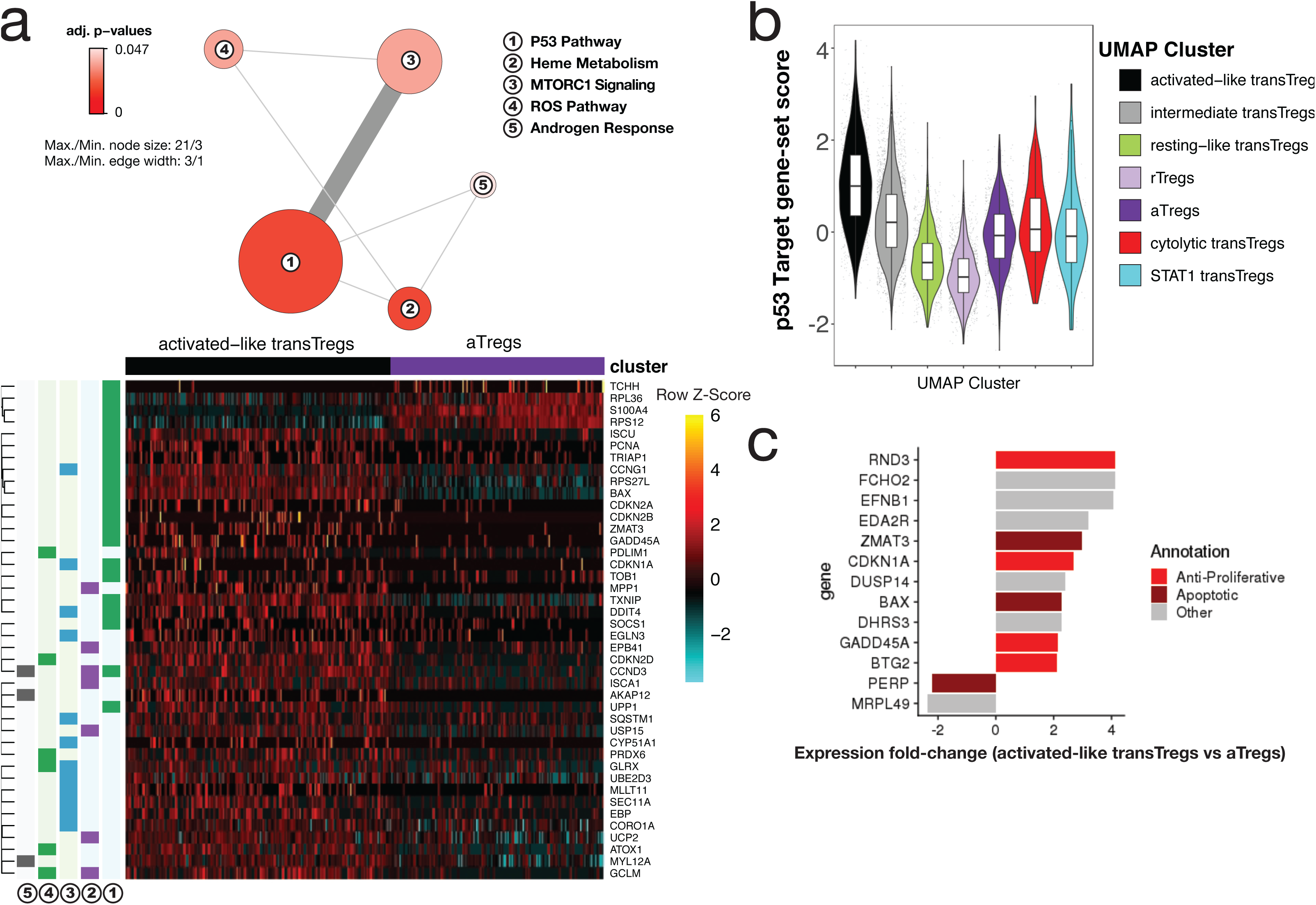
Highly activated transferred Tregs express high levels anti-proliferative and apoptotic genes. a) Network diagram of pathway terms enriched in DE gene list up in activated-like transferred Tregs vs aTregs (top). Expression values of genes from pathways in (a) in activated-like transferred Tregs and aTregs (bottom). b) p53 target gene-set scores by UMAP clusters from Fig. 4b. c) Top p53 target genes DE between activated-like transferred Tregs and aTregs manually annotated as either Anti-proliferative, Apoptotic, Proliferative, Anti-apoptotic or Other.

## DISCUSSION

In this study, we demonstrated that in NHP, the combination of rapa with low-dose IL2 significantly improves persistence of transferred Treg compared to rapa alone, effectively increasing adoptively transferred Treg half-life from 2.7 to 19 days across a large Treg dose range. We then used single cell transcriptomics to show that FOXP3- expressing, endogenous Tregs self-aggregate into two predominant groups of cells that exhibited gene expression patterns consistent with either resting- or activated-Tregs. We further show that early after transfer, circulating ex-vivo expanded Tregs display heterogeneity with respect to resting- and activated-Treg gene expression. Most importantly, we found that cells with the highest activated-related Treg gene expression levels also expressed high p53 target gene levels including those predominantly associated with apoptosis. By day 20, detectible circulating Tregs became uniform in rTreg related gene expression and exhibited far lower levels of p53 target gene expression.

While these data provide rationale for employing IL2+rapa as supportive therapy for adoptively transferred Tregs, they also raise important questions. Although we found that many activated-like Tregs express apoptotic genes after adoptive transfer, it is not yet clear if a subset of transferred activated Tregs has the capacity to transition to longer-lived cells associated with a resting-like gene expression profile. One study of expanded activated CD8 cells derived from central memory (but not effector-memory) CD8 cells found that some of the transferred T effectors re-acquired central memory markers including CD62L as late as 56 days after transfer^46, 47^. Although it is controversial whether Tregs have differentiation states that parallel Tconv cells^48^, if Tregs also have this capacity for central memory differentiation, it is possible that some or all of the resting-like transferred cells were, at one point, aTregs.

Regardless of the origin of these resting-like transferred cells, these cells clearly persist in the setting of IL2+rapa. It has been shown previously that expanded human activated memory (CD62L-CD45RO+) CD8 subsets exhibit higher sensitivity to cytokine (IL2) withdrawal-induced cell death (CWID) compared to their central memory (CD62L+CD45RO+) counterparts^31^. As high IL2 levels are required to expand primate Tregs ex-vivo, it is likely that such Tregs would be particularly susceptible to IL2-CWID. In support of this hypothesis, a study found that the pro-apoptotic protein BAX was upregulated after IL2 withdrawal in cultured murine Tregs and Treg apoptosis could be mitigated by a p53 inhibitor and enhanced in transgenic BAX-expressing Tregs^49^. Our findings of specific and near uniform upregulation of BAX in activated-like transferred Tregs and high levels p53 target gene expression is consistent with these findings. Placing our data in the context of both prior studies, it is likely that adjunctive, low-dose IL2, in the presence of rapa, supports the persistence of cells with a more resting-like gene expression profile. Importantly, multiple studies have shown the immunosuppressive benefit of CD62L+ Tregs over CD62L(-) Tregs^13, 48, 50–53^, further pointing to the use of IL2+rapa as an adjunctive strategy to favor or enhance CD62L+ Tregs for clinical benefit.

Throughout this work we also identified a small number of previously-undescribed Treg cells expressing high mRNA levels for STAT1. Moreover, we found evidence for these cells in prior scRNA-seq studies of murine nonlymphoid^41^ Tregs, confirming their existence across species. These STAT1-Tregs cluster with other STAT1 expressing CD4 T cells and although it is not yet clear what expression of STAT1 targets may functionally represent in Tregs, it has been shown that both the abundance and absence of STAT1 signals are critical to Treg function. Mice with Tregs deficient in STAT1 exhibit an increase in development of experimental autoimmune encephalitis^54^, while increased STAT1 signaling in Tregs has been associated with aberrant Treg homeostasis in patients with systemic lupus erythematosis^55^. Furthermore, dominant gain-of-function STAT1 mutations have been described in patients with an immune dysregulation-polyendocrinopathy-enteropathy-X-linked-like syndrome but with wild-type FOXP3^56^. Finally, in mice, autocrine signaling of interferon-gamma in Tregs has been shown to augment the tolerance to allograft^57^. Further clarification on the role of STAT1 and STAT1 target genes in Tregs is clearly needed.

This study fits within a larger body of work aiming to identify the most ideal adjunctive treatment strategy to pair with ex-vivo expanded, adoptively transferred Tregs. Choosing the appropriate adjunctive immunosuppressant is especially critical given that previous studies have suggested that it is likely that cellular strategies may not be able to sufficiently control inflammation alone and adjunctive agents will be an important pillar of disease control, tolerance induction, or allograft acceptance^3^. As many patients who could theoretically benefit from ex-vivo expanded Tregs will be receiving immunosuppressants and would likely need to continue these interventions at the time of cell infusion. Choosing the preferred immunosuppressant to synergize with and not mitigate the suppressive function is important in optimizing adoptively transferred Treg efficacy. Studies comparing adjunctive therapies for ex-vivo expanded Tregs have been particularly challenging because historically such transferred cells have limited persistence and circulating cells have been insufficient for functional analyses. Here, we provide a technical framework for evaluating adjunctive immunosuppression for ex-vivo expanded, adoptively transferred Tregs that uses a combination of multiparameter flow-cytometry and single cell transcriptional profiling to study the cells longitudinally after adoptive transfer.

In summary, we show that IL2+rapa prolongs adoptively transferred Treg persistence and results in stable expression of transcripts associated with Treg identity and function. Lastly, these data are the first to employ a single cell transcriptomic study of adoptively transferred Tregs in a large animal model and highlight the power of this technology to more precisely inform adoptive Treg immunotherapy.

## Acknowledgement

SNF is supported by an ACS Mentored Scholar award. This work was funded by the following NIH grants: R01 HL11879, R01 AI 34495, 2PO1 CA065493 to BRB and R01 HL095791, U19 HL129902 to LSK. BRB receives remuneration as an advisor to Kamon Pharmaceuticals, Inc, Five Prime Therapeutics Inc, Regeneron Pharmaceuticals, Magenta Therapeutics and BlueRock Therapeuetics; research support from Fate Therapeutics, RXi Pharmaceuticals, Alpine Immune Sciences, Inc, Abbvie Inc., Leukemia and Lymphoma Society, Childrens’ Cancer Research Fund, KidsFirst Fund and is a co-founder of Tmunity

## Author Contributions

SNF and LSK designed this study with input from BRB. SNF, CL, VT, DH, and KMB performed experiments. SNF performed analysis of data with input from all authors. SNF, LSK and BRB wrote manuscript with input from all authors.

## Notes

#### Summary of Updates

Including all authors using web portal

https://dataview.ncbi.nlm.nih.gov/object/PRJNA577285

