## Supplementary Material for "IL2 enhances ex-vivo expanded regulatory T cell persistence after adoptive transfer"

SUPPLEMENTARY INFORMATION

Supplementary Table 1

Supplementary Table 2

Supplementary Figures 1-6

Expanded Materials/Methods

Supplementary References

SUPPLEMENTARY TABLE AND FIGURE LEGENDS

Supplementary Table 1 – Doses of adoptively transferred ex-vivo expanded Tregs.

Supplementary Table 2 – GSEA Leading Edge genes from Figure 3e.

Supplementary Figure 1 - a) Strategy used to sort populations for downstream scRNA-seq. Lymphocytes were first identified by light scatter, then by expression of CD3 and CD4. CD3+CD4+CFSE+ cells were sorted (green), while CFSE- cells were further sorted based on their expression of CD25 and CD127.b) Representative plot of CD25 and CD127 expression in CFSE+ cells.

Supplementary Figure 2 - **Transferred Tregs do not have a CFSE dilutional pattern or expression profile consistent with proliferation after adoptive transfer.** a) CFSE labelling of CD4+ lymphocytes positive for CD25 and FOXP3 (Tregs) for animal R.401 are shown at various timepoints after adoptive transfer. Mean fluorescence intensity of CFSE positive cells shows kinetics that are not consistent with ongoing proliferation of cells. b) Single cell expression profiles of clusters from Figure 3b indicate near absent transcript of the proliferation marker Ki-67 and CCNB2 in cells from tSNE clusters C1 and C2.

Supplementary Figure 3 – **Cells from clusters low in SELL expression, express high levels of the memory T cell genes, CD58, CD63, LGALS3 and S100A4.** Violin plots showing log-normalized UMI counts of memory T cell genes across tSNE clusters in Figure 3a.

Supplementary Figure 4 - **Cells sorted from endogenous Treg gate (CD3+CD4+CD25hiCD127loCFSE-) exhibit similar gene expression patterns to published profiles of sorted CD25hiCD4 T cells and reveal small subset of ISG15, MX1 high cells that express FOXP3.** a) tSNE clustering of endogenous NHP Tregs colored by unsupervised louvain clustering (left); tSNE clusters in (a) scored for expression of published human gene signatures of rTregs, aTregs and Fr.III cells. b) tSNE embedding in (a) colored by expression of gene.

Supplementary Figure 5 – **Tissue resident Tregs (GSE109742) show clusters of cells expressing high levels of STAT1 targets.** a) Cells from GSE109742 samples were downloaded, normalized and dimensionality reduction was performed. b) Unsupervised louvain clustering was performed on PCs obtaining clusters that largetly identified Treg and Tconv clusters in addition to two other groups: 1) ‘Stat1’-Tregs and 2) ‘Other’. Cells in Stat1 cluster were enriched for expression of Mx1 and Isg15 (c,d), Ifnar1 (d) and still expressed canonical Treg markers (Foxp3, il2ra) (d).

EXPANDED MATERIALS AND METHODS

**NHP Ethics Statement:** This study used specific pathogen-free, juvenile rhesus macaques that were housed at the Yerkes National Primate Research Center and the Washington National Primate Research Center. The study was conducted in strict accordance with USDA regulations and the recommendations in the Guide for the Care and Use of Laboratory Animals of the National Institutes of Health. They were approved by the Emory University, University of Washington, and Seattle Children’s Research Institute’s Institutional Animal Care and Use Committees.

**Isolation and ex-vivo expansion of Tregs:** CD4^+^CD25hiCD127low Tregs, from autologous donors were aseptically obtained using our previously described protocol^1^. In brief, Tregs were flow-sorted from CD4 T cells in peripheral blood samples after Ficoll purification and immunomagnetic enrichment for CD4, and were subsequently stimulated with anti-rhesus-CD3 and anti-human CD28 coated microbeads, such that Tregs were stimulated and re-stimulated on days 0, 12 and 24 post-purification (**Figure 1a**). As previously described^2^, on day +34, cells were treated with rapamycin and on day +36 were washed free of rapamycin and magnetic beads were removed. The Treg phenotype in the expanded cells after adoptive transfer was assessed by staining for CD3 (clone SP34-2, BD, San Jose, CA), CD4 (clone SK3, BD), CD25 (clone 4E3, Miltenyi Biotec), CD127 (clone eBioRDR5, eBioscience, San Diego, CA) and FoxP3 (clone 150D, BioLegend) using the FoxP3 Fix/Perm Buffer Set (BioLegend, San Diego, CA). Data were acquired on a Fortessa flow cytometer and analyzed using FlowJo software (Treestar, Ashland, OR). Positively stained cells were identified using appropriate isotype-control antibodies. Treg batches were tested for sterility (University Washington Medical Center, Clinical Microbiology laboratories) prior to adoptive transfer.

**Infusion and tracking of CFSE labeled Tregs:** Tregs were labeled with 10μM of CFSE (Invitrogen, Grand Island, NY), washed free of excess CFSE and subsequently cultured at 37^o^C in a CO_2_ incubator in X-Vivo-15 media supplemented as previously described^2^ for 24h prior to infusion into autologous recipients. This 24 hour culture period allowed CFSE toxicity to be minimized, as previously described^3-5^. Just prior to infusion, the CFSE-labeled cells were washed three times with normal saline supplemented with 2% autologous serum, and passed through a 70 μm filter to prevent cell clumping. Their viability and cell number were determined by trypan-blue staining, and the cells were resuspended at 20x10^6^ live cells/ml and immediately infused into recipients at Treg doses ranging from 3.1x10^6^-30.4x10^6^/kg. The analysis of Treg persistence was performed using baseline cytometric measurement of transferred (CFSE^+^) Tregs was made within 60 minutes of this infusion by identifying the percentage of CD3^+^CD4^+^CD25^+^FoxP3^+^ cells that were CFSE+. All subsequent measurements of transferred Tregs (CD3^+^CD4^+^CD25^+^FoxP3^+^CFSE+) were expressed in terms of a percent relative normalized to the “time 0” value.

**Immunosuppression and IL2:** Animals were treated with rapamycin (LC laboratories) daily by intramuscular injection, which was begun two weeks before the Treg infusion, to achieve a target trough level of 5-15 ng/ml assayed from whole blood using liquid chromatography-tandem mass spectrometry after protein precipitation, and continued for the length of longitudinal analysis. Aldesleukin, referred to herein as IL2, (trade-name – ‘Proleukin’ Prometheus) was reconstituted according to the manufacturer’s instructions and given daily by subcutaneous route at a dose of 1 million units/m2 starting 5 days before adoptive transfer until day 60 post-transfer.

**Single cell RNA-seq library prep:** Cryopreserved PBMCs from animal R.401 were recovered and stained with surface antibodies (Supplementary Figure 1). Single cell libraries from sorted populations were generated using a UMI-based droplet-partitioning platform (10X Genomics) and sequenced using a NextSeq 500 (Illumina). Resulting reads were processed using Cellranger software (10X Genomics) and downstream analysis was performed using Monocle^6,7^ using a negative binomial model of distribution with fixed variance. Analysis was limited to those genes that had a detectible UMI or more in at least 1 cell. Scripts detailing the workflow have been uploaded to the GitHub repository (www.github.org/scfurl/NHPTregIL2) and all sequence data have been uploaded to the NIH SRA and GEO (PRJNA577285).

**Single cell RNA-seq QC:** Cells whose UMI counts for mitochondrial genes greater than 5% were excluded from downstream analysis. Cells with between 500 and 2000 genes expressed per cell were included in the analysis.

**Derivation of gene-sets, GSEA and single cell gene-set scores:** The Treg-specific gene-set (**Figure 3d**) was derived from GSE90600 expression data. Read counts were obtained from GEO and analyzed using DESeq2^8^. Differentially expressed genes across cell types were calculated using thresholds of a DESeq2-calculated and adjusted p value of < 0.05 in addition to a log-fold change cutoff of > 1.5. Non-human primate rTreg and aTreg gene-sets were derived from differentially expressed genes between clusters C3 and C4 using Monocle^6^. Gene-sets for p53 targets were taken from a published meta-analysis of p53 target genes^9^. Gene set enrichment analysis (GSEA) on single cell data expression data was performed by first: Ranking gene expression between two groups of cells using the full model coefficient derived from the ‘differential gene test’ function in Monocle. Secondly, these ranked lists were interrogated for enrichment (using the Piano^10^ package) for any given gene-set. Single cell gene-set scores were calculated so that for any given cell, a centered and scaled the average normalized pseudocount for all genes in the geneset.

**Clustering, Differential Expression and Enrichment Analysis:** All clustering was performed using Monocle. For unsupervised clustering of cell expression profiles, expression matrices from Cellranger were first normalized by size factors and dispersion. Genes to use for clustering were then selected based on whether they met on thresholds for mean expression and dispersion. Principal components analysis (PCA) was then performed on the expression matrix of the selected genes. The resulting PCA matrix was then subsetted to exclude those PCs that contributed little to the overall variance. Dimensionality reduction using tSNE^11^ or UMAP^12^ was then performed on the subsetted PCA matrix. As stated above, DE was performed in Monocle using a conservative significance threshold (FDR of 0.01). Enrichment analysis was performed using the Piano software package^10^ and enrichment terms from the MSigDB^13^. Functional gene annotations were taken from GeneCards^14^ unless otherwise specified.

**Published datasets:** Datasets (GSE90600^15^ and GSE109742^16^) were downloaded for GEO. GSE90600 processed gene counts were used to generate DE genelists between rTregs, eTregs, FrIII cells and RA+ and RA- Tconv using DESeq2 with significance thresholds of 0.05 (after correcting for multiple hypotheses). Processed read counts for samples in GSE109742 were downloaded and samples from muscle, spleen and visceral adipose tissue (VAT) were analyzed using monocle. Clustering was performed as above.
