## Supplementary figures and images for "IL2 enhances ex-vivo expanded regulatory T cell persistence after adoptive transfer"

### Supplementary Figure 1

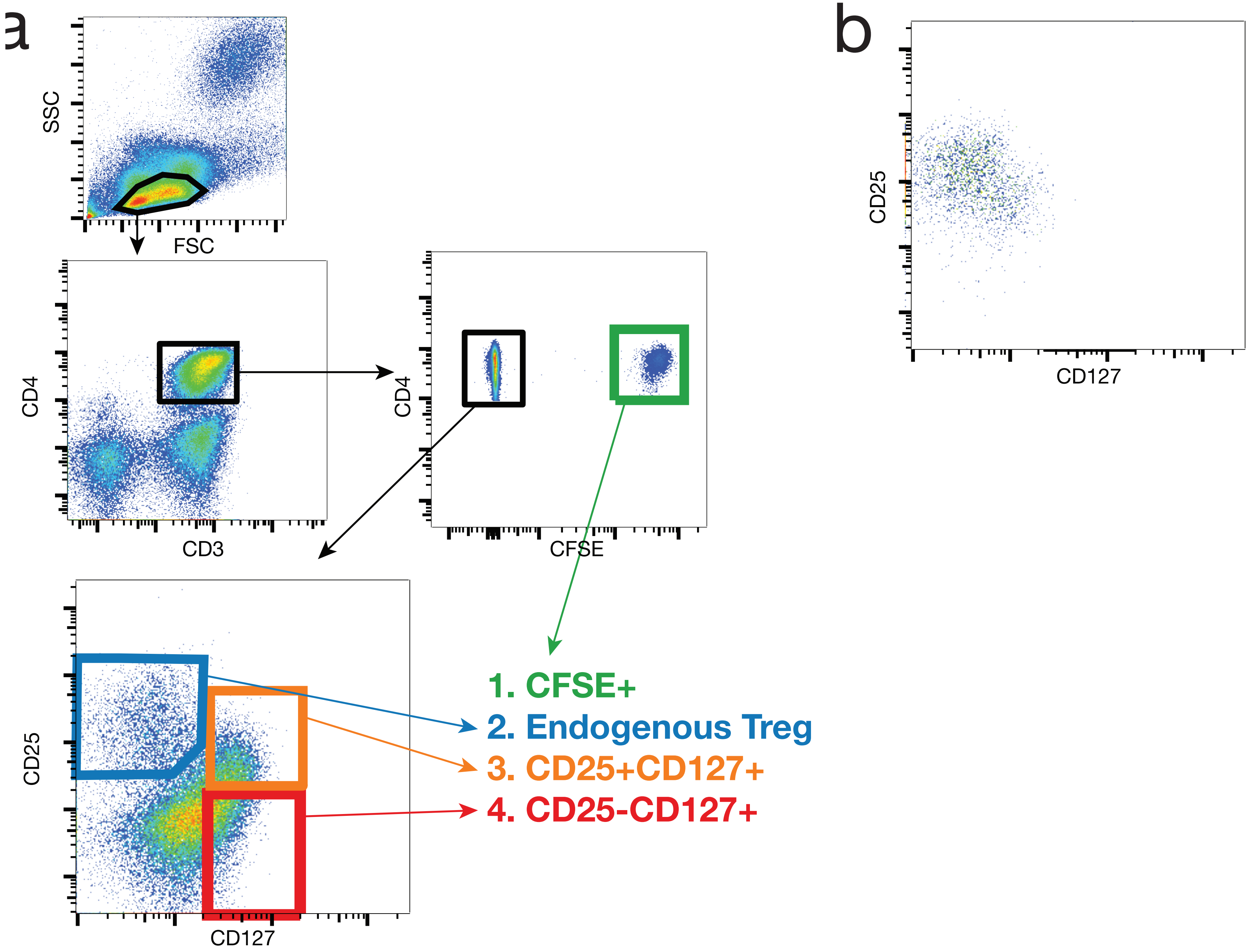


SUPPLEMENTARY FIGURE 1

### Supplementary Figure 2

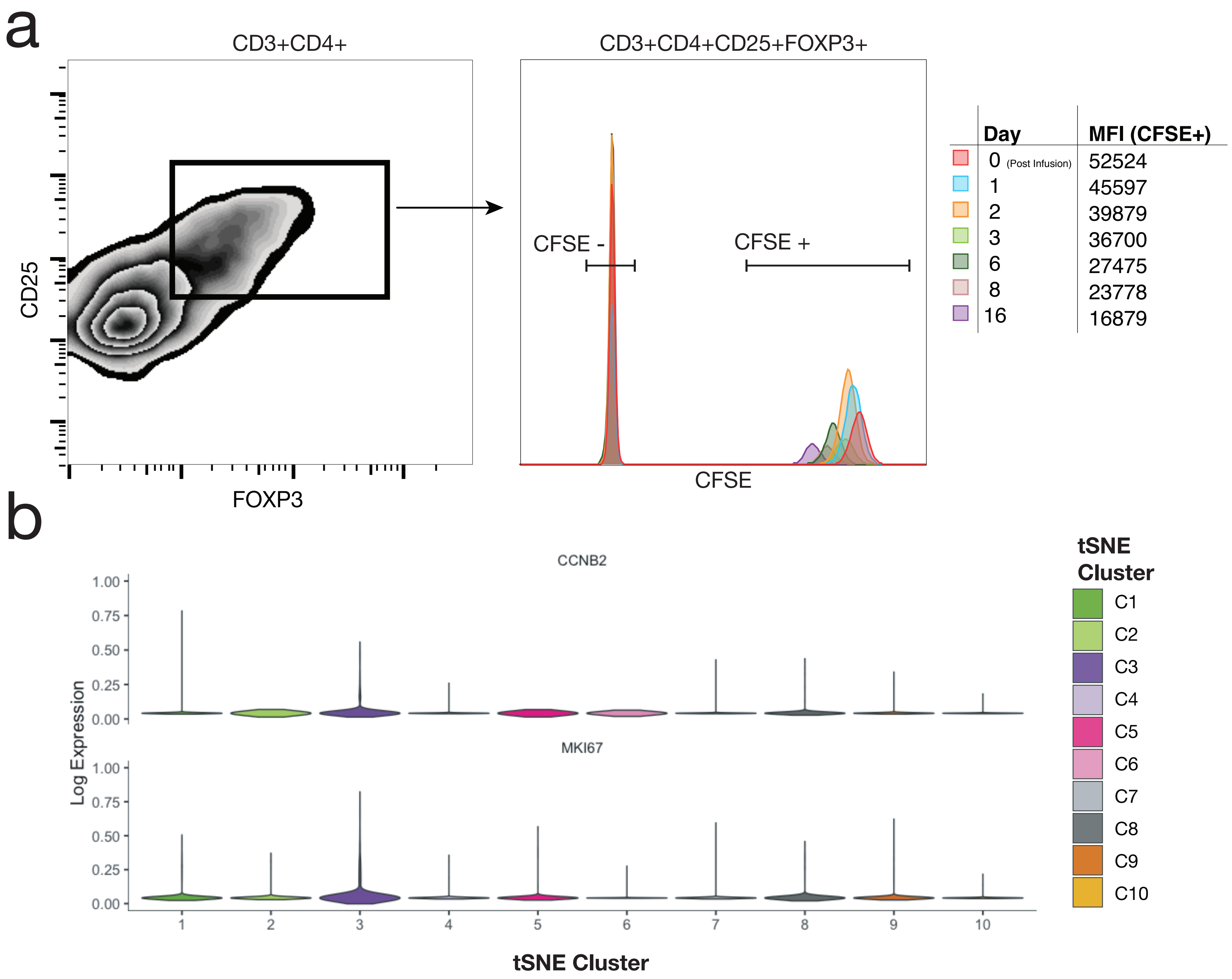


SUPPLEMENTARY FIGURE 2

### Supplementary Figure 3

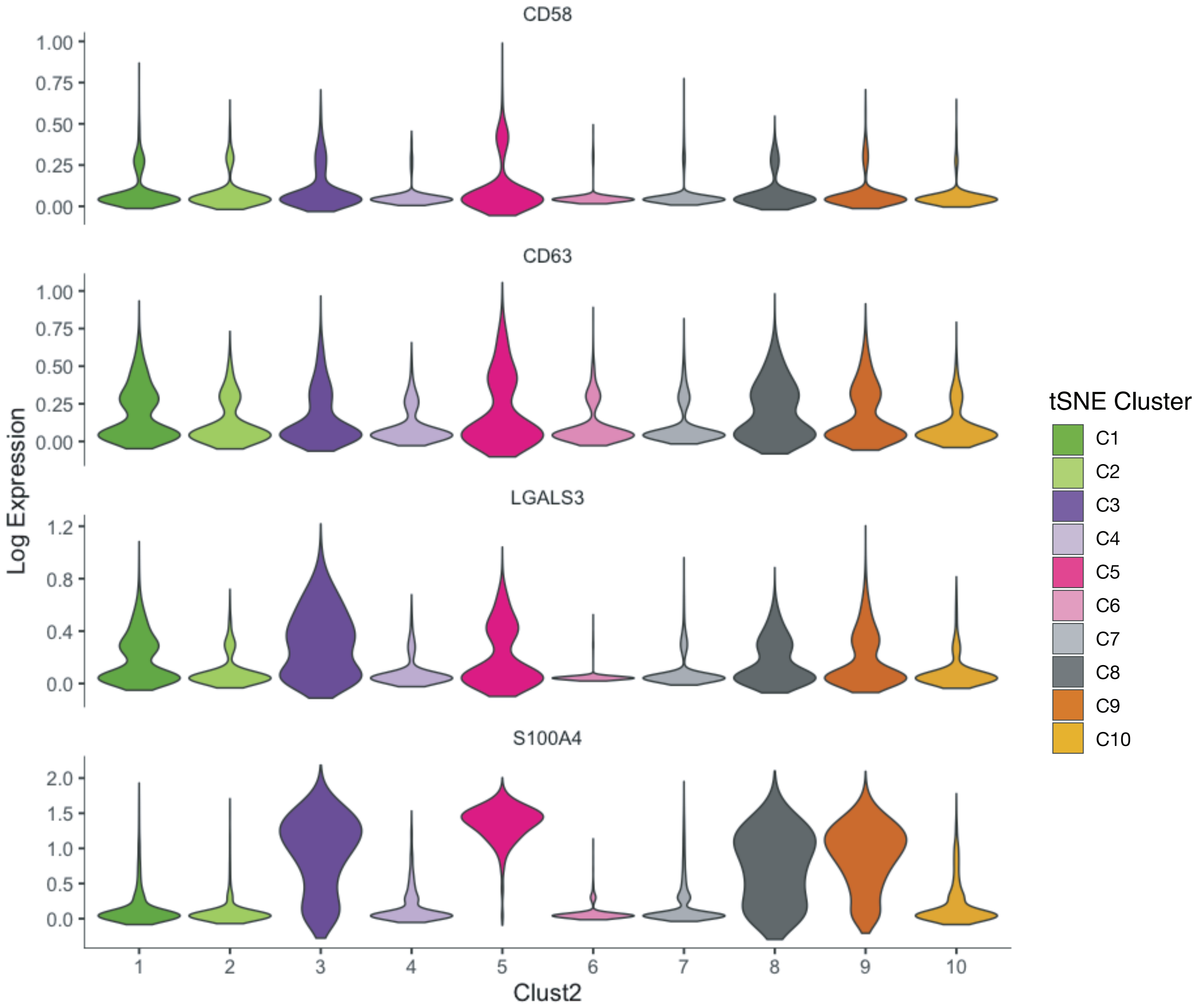


SUPPLEMENTARY FIGURE 3

### Supplementary Figure 4

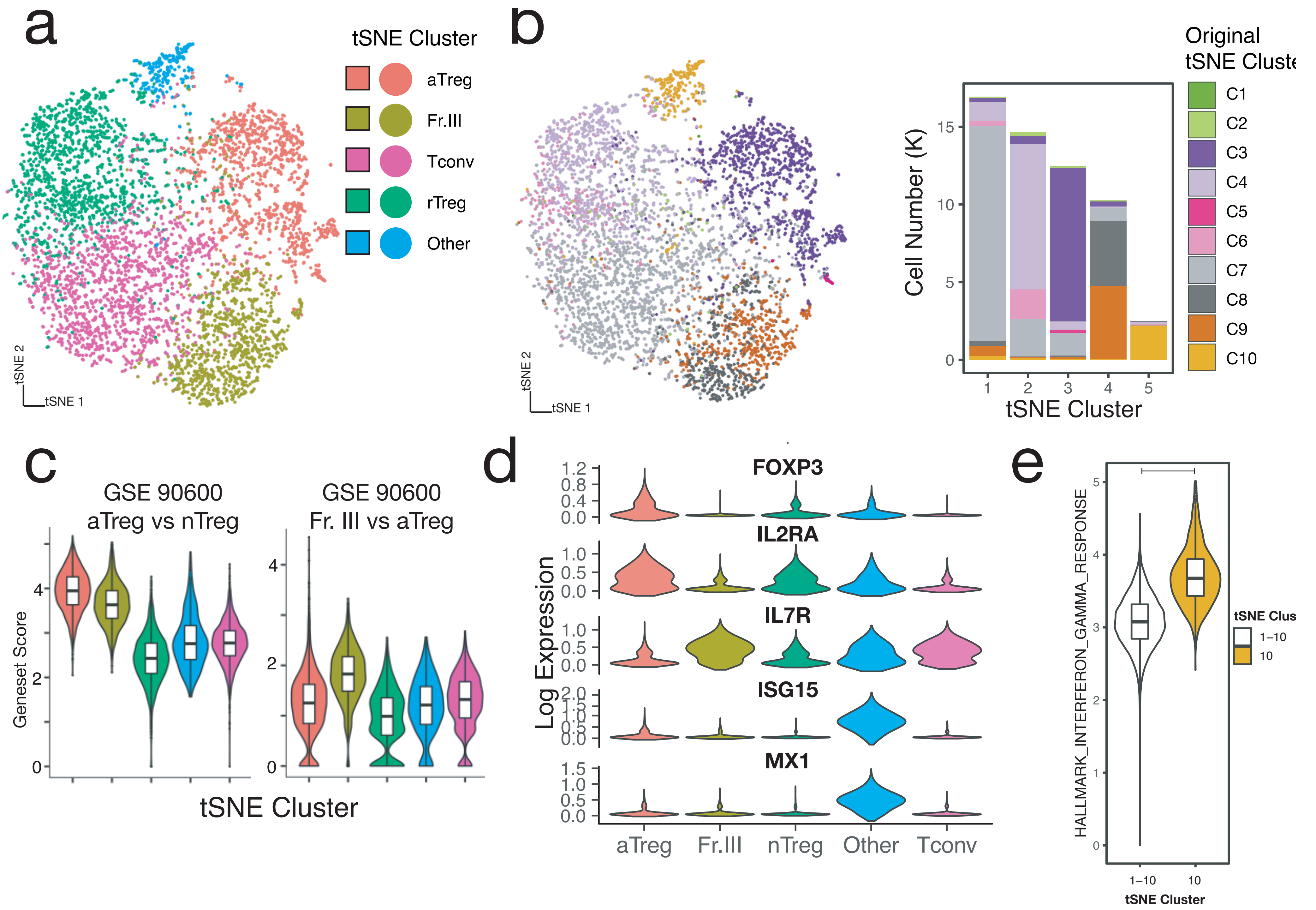


SUPPLEMENTARY FIGURE 4

### Supplementary Figure 5

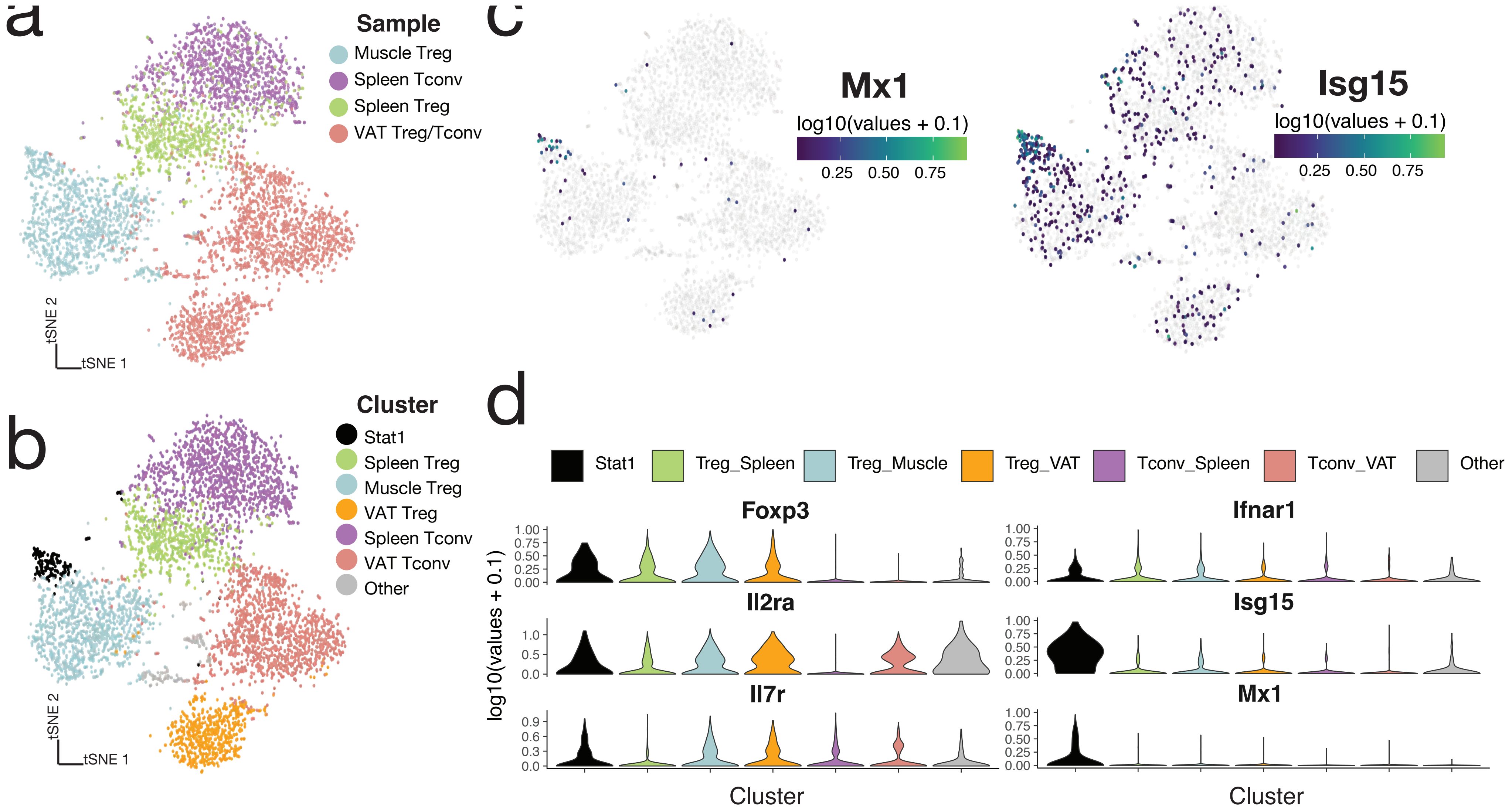


SUPPLEMENTARY FIGURE 5
