## Supplementary Table 1 for "IL2 enhances ex-vivo expanded regulatory T cell persistence after adoptive transfer"

| Animal ID | Total Live Cells Infused | Weight (kg) | Cell Dose (Cells / kg recipient) |
|---|---|---|---|
| R.401 | 4.27E+08 | 14.7 | 2.90E+07 |
| R.402 | 4.80E+08 | 15.8 | 3.04E+07 |
| R.403 | 2.50E+07 | 8.2 | 3.10E+06 |

SUPPLEMENTARY TABLE 1
